# No support for cell size as a driver of tissue-level metabolic rates at the upper limits of animal cell size

**DOI:** 10.64898/2026.06.17.733039

**Authors:** Michael W. Itgen, Adam J. Chicco, Rachel Lockridge Mueller

## Abstract

Evolutionary diversity in metabolic rate underlies differences in physiology, morphology, and life history across the tree of life. Cell size has been proposed as an important determinant of metabolic rate. The mechanisms underlying this proposed connection are based on the lower surface area to volume ratios in larger cells. As relative surface area decreases, the cost of maintaining ion gradients across the cell membrane through action of the Na+/K+-ATPase pump are posited to decrease, lowering overall metabolic costs. Despite strong theoretical support for this model, and its incorporation into broader models of life history evolution, empirical measurement of Na+/K+-ATPase activity in species that differ in cell size has been lacking. Here, we study nine species of salamanders of the genus *Plethodon* that span a large range of cell sizes approaching the animal upper limit. We compare basal cellular respiration rates, relative cost of the Na+/K+-ATPase pump, and maximal mitochondrial respiration rates in liver and heart tissue. Contrary to predictions, we find no support for a relationship between cell size and any of these mitochondrial respiratory variables. We reconcile this surprising result with broader phylogenetic studies showing a lack of correlation between cell size and metabolic rate at the organismal level.

## Introduction

Metabolic rate is the rate of total energy expenditure by an organism to sustain all of the biochemical processes underlying life. Across taxa, metabolic rate varies dramatically, associated with differences in development, physiology, morphology, and life history (Gillooly, et al. 2001; Brown, et al. 2004; Killen, et al. 2016). Body size is a strong predictor of metabolic rate, and diverse causal mechanisms contribute to the allometric scaling relationship between these two variables (Kleiber 1947; Glazier 2005; Kozłowski, et al. 2020). Taxa of the same body size can also differ substantially in metabolic rate, in some cases driven by the same causal mechanisms that underlie allometric scaling with body size (Makarieva, et al. 2008; White and Kearney 2013).

Differences in cell size have been proposed to drive differences in metabolic rate, both associated with and independent of body size (Glazier 2022). As cells increase in size, assuming their shape remains constant, the cell surface area-to-volume ratio (SA/V) decreases (Chan and Marshall 2010; Marshall, et al. 2012). Thus, the relative amount of cell (i.e. plasma) membrane and membrane-associated proteins decreases, and the relative amount of cytoplasm increases (Miettinen, et al. 2017). Decreased SA/V in larger cells can affect metabolic rate in two ways that are not mutually exclusive − by decreasing metabolic demand and by limiting metabolic supply. More specifically, larger cells have been hypothesized to require less energy for maintainance of the ion gradients across the cell membrane that create the membrane potential (Davison 1955; Kozowski, et al. 2003). Na+ and K+ ions are pumped across the cell membrane against their concentration gradients by Na+/K+-ATPase enzyme activity (Skou 1957; Fedosova, et al. 2022). In larger cells, the lower SA/V ratio has been posited to result in less passive diffusion of these ions across the cell membrane, thus requiring less energy expenditure by the cells for their active transport by Na+/K+-ATPase, resulting in decreased metabolic demand (Johnston, et al. 2006). Diffusion also limits metabolic supply in larger cells, as increased intracellular distances and decreased membrane surface area constrain the efficacy of diffusion of oxygen and metabolites (Egginton, et al. 2002; Miettinen and Björklund 2017). Cell size is strongly positively correlated with genome size, as DNA content is a partial determinant of cytoplasmic volume (Gregory 2005; D’Ario, et al. 2021). Because it is much more easily measured, genome size is often used as a proxy for cell size in studies that examine the relationship between cell size and metabolic rate (Uyeda, et al. 2017; Gardner, et al. 2020).

Cell size and its putative effects on metabolic rate have been incorporated into more general conceptual frameworks for life history evolution, specifically the idea that life history strategies fall on a continuum from “frugal” to “wasteful” (Szarski 1983). Frugal strategies encompass organisms with low metabolic rates, the capacity to inhabit oxygen- or resource-limited environments, low levels of activity, and slow rates of growth and reproduction. Wasteful strategies encompass organisms with high metabolic rates, the capacity to rapidly exploit available resources, high levels of activity, and rapid rates of growth and reproduction (Szarski 1983). In its original conception, the frugal strategy was typified by the salamanders and the lungfishes, both of which have gigantic cells and genomes (∼9 Gb – 120 Gb), and the wasteful strategy was typified by birds, which have small cells and genomes (0.9 Gb – 1.3 Gb) (Szarski 1983; Zhang, et al. 2014; Schartl, et al. 2024; Gregory 2025). Large cell size was interpreted as a target of natural selection for “economy of resources” (i.e. low metabolic demand), and small cell size for high metabolic capacity and fast development (Szarski 1983).

Since this original conception, additional hypotheses have expanded on this idea, including the “Theory of optimal cell size (TOCS)” and the “optimal fiber number hypothesis,” both of which invoke cell size as a target of natural selection, optimized to balance the competing forces of minimizing ion gradient maintenance costs (demand) and allowing diffusion of oxygen and metabolites (supply) (Johnston, et al. 2003; Johnston, et al. 2006; Schramm, et al. 2021). Comparative studies from diverse taxa including fishes, birds, lizards, molluscs, beetles, ants, and crustaceans have found results consistent with these ideas. In fishes, species with lower diffusional constraints associated with lower metabolic rates or higher hemoglobin oxygen binding affinities have larger muscle cell diameters (Johnston, et al. 2006). In galliform birds and carabid beetles, species or sexes with larger body sizes and lower mass-specific metabolic rates have larger cell sizes (Czarnoleski, et al. 2018; Schramm, et al. 2021). In geckos and ants, species in which cell size increases throughout ontogeny show sublinear scaling of metabolic rate with body size (Chown, et al. 2007; Starostová, et al. 2013). In mussels, individuals living high on the shores in the rocky intertidal, where access to food resources is limited, have larger muscle cell diameters and lower aerobic capacity than conspecifics in more resource-rich microenvironments (Dowd and Jimenez 2019). In seeming contrast to all of these taxonomically shallow comparisons, however, phylogenetic tests for the predicted negative relationship between genome size (proxy for cell size) and metabolic rate within and among all of the extant vertebrate clades fail to identify genome/cell size as a significant predictor of metabolic rate (Uyeda, et al. 2017; Gardner, et al. 2020). Taken together, these results suggest that, despite strong theoretical predictions and some empirical support among closely related taxa, cell size is not a universal predictor of metabolic rate. These conflicting bodies of evidence reveal a need for experimental approaches that directly measure ion gradient maintenance costs in cells of different sizes and their effects on metabolic rate.

Despite this clear need, however, very few studies have measured the cost of Na+/K+-ATPase enzyme activity, and the percentage of total energy expenditure it accounts for, across cells of different sizes. Jimenez et al. (2013) compared large versus small muscle cells, which were associated with large versus small body size, in 16 species of crustaceans and fishes. For each size class in each species, they measured the total and fractional cost of Na+/K+-ATPase activity, revealing that the cost of ion gradient maintenance decreases as a function of decreasing SA/V ratio in larger cells (Jimenez, et al. 2013). Cadart et al (2023) used the model frog genus *Xenopus*, in which ploidy increases can be experimentally induced, to compare total energy expenditure and Na+/K+-ATPase activity cost between diploid and triploid embryos. Because cell size and DNA content are correlated, triploid embryos have larger cells; as predicted, triploids have lower metabolic rates than diploids because of lower ion gradient maintenance costs (Cadart, et al. 2023). Additional such experimental studies that include: 1) multi-species comparisons in which cell size is variable, but body size and ploidy are invariable, 2) cell types that encompass a range of morphologies and functions (muscle cells are unusual in being multinucleated, cylindrical, and excitable); and 3) more of the natural diversity in cell size found across the tree of life are necessary to quantify the metabolic impact of cell size and integrate its role with other known drivers of metabolic rate (e.g. relative sizes of metabolically intensive organs, membrane composition, mitochondrial proton leak rates, protein turnover rates, organellar density) (Welle and Nair 1990; Turner, et al. 2005; White and Kearney 2013).

The salamander genus *Plethodon* meets these requirements for a model taxon for cell size research (Itgen, et al. 2022). *Plethodon* encompasses 58 named species of diploid, lungless, direct-developing, terrestrial salamanders with a broad range of genome sizes (23–67 Gb) and cell sizes shaped by stochastic evolutionary processes, but uniformity or lower diversity in potentially confounding variables (e.g., life history, body size) that also impact metabolic rate (Petranka 1998; Newman, et al. 2016; Itgen, et al. 2022; Mueller, et al. 2023) (Table 1). The two main clades within *Plethodon* – the eastern and western clades – diverged ∼45 mya (Kumar, et al. 2017), but evolutionary stasis has been so pervasive throughout the group’s history that molecular data more than tripled the number of named species over the last several decades (Highton 1995; AmphibiaWeb 2025).

**Table 1.** Mean ± standard deviation for genome size, cell size, and body size of focal taxa from Itgen et al. (2022).

| Species | Genome size<br>(Gb) | Hepatocyte Area<br>(µm<sup>2</sup>) | Snout-vent length |
| --- | --- | --- | --- |
|  |  |  | (mm) |
| *Plethodon cinereus* | 29.3 ± 1.34 | 373.8 ± 59.5 | 40.1 |
| *P. montanus* | 36.0 ± 3.4 | 515.3 ± 34.8 | 49.7 |
| *P. cylindraceus* | 37.1 ± 5.4 | 550.4 ± 27.2 | 70.4 |
| *P. metcalfi* | 38.3 ± 1.04 | 492.7 ± 90.3 | 61.9 |
| *P. glutinosus* | 38.9 ± 3.94 | 498.6 ± 42.9 | 66.9 |
| *P. vehiculum* | 46.4 ± 7.08 | 740.1 ± 71.4 | 50.5 |
| *P. dunni* | 52.3 ± 4.06 | 755.2 ± 108.6 | 68.7 |
| *P. vandykei* | 54.6 ± 2.2 | 943 ± 227.5 | 54.2 |
| *P. idahoensis* | 67.0 ± 2.31 | 984.5 ± 371 | 58.1 |

Here, we analyze nine species from the salamander genus *Plethodon* that span the full range of genome/cell sizes. We test the hypotheses that 1) larger cell sizes are associated with lower basal rates of cellular respiration, 2) lower basal rates of cellular respiration in larger cells reflect lower costs of maintaining ion gradients across the plasma membrane through Na+/K+-ATPase activity, and 3) larger cells have lower maximum mitochondrial respiratory capacity. We test all three of these hypotheses using liver tissue (primarily composed of hepatocytes), and we include heart tissue (cardiomyocytes) as an additional test of Hypothesis 3. In contrast to strong theoretical predictions, our results reveal that evolutionary increases in cell size across *Plethodon* are not associated with decreased basal rates of mass-specific, tissue-level cellular respiration, decreased costs of ion gradient maintenance costs, or decreased maximum respiratory capacity. We re-examine the classic evolutionary scenario connecting cell size, cell SA/V ratio, membrane potential maintenance cost, and selection on metabolic demand in light of this surprising result, and we emphasize the importance of empirical tests using a wide range of tissues and species in evaluating theoretical models to explain broad evolutionary patterns.

## Methods

### Animal and tissue collection

We field-collected 5 adult individuals of *Plethodon cinereus*, *P. cylindraceus*, *P. dunni*, *P. glutinosus*, *P. idahoensis*, *P. metcalfi*, *P. montanus*, *P. vandykei*, and *P. vehiculum*. Genome sizes, hepatocyte cell sizes, and body sizes for all species are summarized in Table 1. All permits were issued to XXXX and the animals were collected between May and August of 2018. *Plethodon idahoensis* was collected from Shoshone county, Idaho, under the wildlife collection permit #180226 issued by the Idaho Department of Fish and Game. *Plethodon cinereus* and *P. glutinosus* were collected from South Cherry Valley and Oneonta, Otsego County, New York, under the New York State Department of Environmental Conservation scientific collection permit #2303. *Plethodon vehiculum*, *P. vandykei*, and *P. dunni* were collected from Pacific County, Washington, under the scientific collection permit # XXXXX issued by the Washington Department of Fish and Wildlife. *Plethodon metcalfi* was collected from Macon County, NC, and *P. montanus* and *P. cylindraceus* were collected from Avery County, NC, under the wildlife collection license # 18-SC01250 issued by the North Carolina Wildlife Resources Commission. Animals were returned to XXXXX in coolers by car and were then kept on wet paper towels in containers at 15°C and were fed fruit flies weekly. Salamanders were euthanized by submersion in buffered 1% MS-222 3 days after last feeding. The protocols for animal research, transportation, husbandry, and euthanasia were approved by the Institutional Animal Care and Use Committee of XXXXX (Protocol 17–7189A).

The livers and hearts were dissected immediately, weighed, and transferred into ice-cold biopsy preservation solution (BIOPS) containing (in mM): 10 Ca-EGTA (0.1 μM free calcium), 20 imidazole, 20 taurine, 50 K-MES, 0.5 dithiothreitol, 6.56 MgCl_2_, 5.77 adenosine triphosphate, and 15 phosphocreatine, pH 7.1. Liver tissue was sectioned into experimental samples with wet weights of ∼15 mg (average = 14.9, range = 7.9-18.7). The heart was dissected out entirely and rinsed of blood; heart fiber bundles were blotted dry on Whatman paper and weighed to achieve sample sizes of ∼2-3 mg (average = 2.2, range = 0.4-5.1). The heart muscle fibers were then gently teased with needle-tip forceps in ice-cold BIOPS followed by incubation with 25 μg/mL saponin in BIOPS on ice for 20 minutes with gentle rocking to permabilize cell membranes. Permeabilized fiber bundles were then transferred to mitochondrial respiration medium (MiR05) containing (in mM) 0.5 EGTA, 3 MgCl_2_ hexahydrate, 60 lactobionic acid, 20 taurine, 10 KH_2_PO_4_, 20 HEPES, 110 sucrose, and 0.1% BSA, pH 7.1 with KOH and kept on ice prior to experimentation.

### Basal cellular respiration rate

Our first goal was to test whether mass-specific, tissue-level metabolic rates differ across cells of different sizes. To this end, basal cellular respiration rates were measured as the rate of oxygen consumption of intact cells per mg of liver tissue. The majority of the cells were expected to be hepatocytes, although we did not identify the percentage of different cell types in the tissue. Basal respiration rates of ∼15 mg of liver tissue were measured using the Oxygraph-2k (O2K) high-resolution respirometer system (Oroboros Instruments GmbH, Innsbruck, Austria) at 25°C in 2 mL of MiR05 in the absence of exogenous substrates (i.e., fueled solely by endogenous substrates present in the tissue from the time of sacrifice). Sample contents are contantly mixed by a stirbar rotating at 750 rpm, which gently disperses cells from liver tissue to reduce substrate and oxygen diffusion limitations on individual hepatocytes. We selected 25°C because Licht and Lowcock (1991) found a significant effect of genome size on metabolic rate in salamanders at this temperature, and although it is above the preferred body temperature, it is still ecologically relevant and below the critical thermal maximum (e.g. for *P. cinereus*, preferred temperature in the laboratory is 16 to 21°C, but the CT_max_ is 32°C for animals acclimated at 15°C) (Feder and Pough 1975; Licht and Lowcock 1991). Experiments were done in a hyper-oxygenated environment (400-450 μM oxygen) by injecting oxygen into the chamber prior to collecting data to avoid any oxygen diffusion limitations on cellular respiration rate (Pesta and Gnaiger 2011; Li Puma, et al. 2020). Mass-corrected oxygen consumption rates (pmol O_2_ /mg tissue wet weight/sec; *^J^*^O^_2_) were monitored in real time by resolving changes in the negative time derivative of the chamber O_2_ concentration.

### Na+/K+-ATPase pump respiratory cost

Our second goal was to test whether larger cells expend less of their total energy budget on Na+/K+-ATPase activity, given their lower SA/V ratio. After obtaining measurements of basal cellular respiration rate (above), we inhibited Na+/K+-ATPase activity by adding a high concentration (60 μM) of ouabain to the samples of intact cells respiring in the Oxygraph chamber, then measuring the associated decrease in respiration rate under the same experimental conditions. We then calculated the proportional change from basal cellular respiration to respiration with inhibited Na+/K+-ATPase to determine its contribution to the basal rate of oxidative metabolism. We confirmed that ouabain had no effect on mitochondrial oxygen consumption rate in permeabilized liver cells or isolated mitochondria in separate preliminary experiments, verifying that the observed lowering of intact-cell respiration rates by ouabain reflects the reduced ATP demand from Na+/K+-ATPase inhibition.

### Mitochondrial maximal OXPHOS-linked respiration rate

Our third goal was to test whether the mitochondria from larger cells have lower maximum respiratory capacity, which we predicted would evolve in response to the lower functional demands on the mitochondria of species with larger cells and lower metabolic rates. We used both liver and heart cells for this experiment; the liver cells were first assessed for basal respiration and Na+/K+-ATPase pump cost (above), but heart cells were not subjected to these initial experiments because of technical limitations. Cell membranes were permeabilized using 25 μg/mL of digitonin while leaving the mitochondrial membrane intact, enabling efficient uptake of respiratory substrates and ADP by mitochondria in the permeabilized cells. Maximum respiratory capacity during oxidative phosphorylation (OXPHOS) was then induced by providing mitochondria with saturating concentrations of substrates to reconstitute tricarboxylic acid cycle flux and maximize supply of electrons to the electron transfer system, including (in mM) 1 malate, 5 pyruvate, 10 glutamate, and 10 succinate in the presence of 2.5 mM of ADP to support maximal ATP synthase activity. 60 μM of blebistatin was added to the heart tissue to inhibit myosin heavy chain activity. Mitochondrial respiration experiments were conducted using the O2K with 2 mL of MiR05 substrate in a hyper-oxygenated environment at 25°C, as above.

### Testing for differences in respiratory phenotypes across cells of different sizes

Having measured respiratory variables across species with different genome/cell sizes (above), our next goal was to test whether these variables (1) differed across species, and (2) correlated with differences in genome/cell sizes, as predicted. We first used a MANOVA to test for among-species differences in: basal cellular respiration rates in liver (largely hepatocytes); Na+/K+-ATPase pump respiratory cost in liver (largely hepatocytes); and maximum oxidative respiratory capacity in mitochondria from both liver (largely hepatocytes) and heart (largely cardiomyocytes). We used follow-up univariate ANOVAs to identify significantly different variables followed by omega-squared tests to estimate effect sizes and post-hoc Tukey tests to identify significantly different species. Statistical analyses were performed in R v 4.6.0 (R Core Team 2025).

We then tested for a correlation between genome size (a proxy for cell size) and these same respiratory variables using phylogenetic generalized least squares (PGLS) models. The phylogeny and genome sizes for our nine focal species of *Plethodon* were taken from Itgen et al. (2022). A Brownian motion model of trait evolution was applied to all variables and included a simultaneous estimation of Pagel’s lambda for each trait (Revell 2010). We performed a separate PGLS for the three liver cell variables (basal cellular respiration rate, Na+/K+-ATPase pump cost, and mitochondrial maximal OXPHOS-linked respiration rate) and heart mitochondrial maximal OXPHOS-linked respiration rate. Genome size was the independent variable in all analyses, and the mean species values for the respiratory traits were the dependent variables. The PGLS analyses were conducted using R v 4.6.0 and the packages *caper* and *ape* (Orme, et al. 2011; R Core Team 2025). As an additional exploratory analysis, we repeated the PGLS for maximal heart mitochondrial respiration rate, omitting one species (*P. montanus*) that was a visual outlier in the PGLS result.

For both ANOVA and PGLS analyses, we excluded several datapoints because their extreme values suggested sample or measurement issues during the experiments: one *P. dunni* sample (Pdunni12) from the Na+/K+-ATPase measurements because the drop in respiration was >2.2-fold higher than the next-highest value; one *P. montanus* sample (Pmont15) from liver maximum mitochondrial respiration rate because estimated levels were >2.2-fold higher than the next-highest value; one *P. montanus* sample (Pmont16) and one *P. vandykei* sample (Pvan7) from heart maximum mitochondrial respiration rate because estimated levels were >2.2-fold smaller than the next-lowest value; and one *P. montanus* sample (Pmont12) from the Na+/K+-ATPase measurements because there was no recorded change in respiration rate following the addition of ouabain, suggesting failed Na+/K+-ATPase inhibition.

### Overall cellular protein concentrations

Given that experimentally induced increases in cell size can result in dilution and/or other changes to the proteome (Neurohr, et al. 2019; Lanz, et al. 2022; Xie, et al. 2022), our final goal was to test whether differences in cell size across species were correlated with differences in overall protein concentrations; because our respiration measurements are standardized by tissue mass, we wanted to ensure that mass was not associated with differences in amounts of protein across species with different cell sizes, which could have impacted metabolic rates independent of ion gradient costs. Protein concentrations of liver tissue samples were measured using the Pierce BCA Protein Assay Kit (Thermo Scientific catalog number 23225) for seven of the nine species using liver samples stored in RNAlater at -80°C: *P. vehiculum, P. cylindraceus, P. dunni, P. montanus, P. metcalfi,* and *P. idahoensis* (3 individuals/species), and *P. glutinosus* (2 individuals). For each sample, the tissue was thawed and dried, and a small subsample was obtained weighing ∼5 mg (3.5 – 6.5). Absorbance of each sample was measured following manufacturer’s protocols. To calculate the percentage of protein per sample from the absorbance, we generated a standard curve with known protein concentrations ranging from 0.03 to 2.0. To test for differences in protein concentration across species, we used an ANOVA. To test for a correlation between genome size (proxy for cell size) and protein concentration, we used PGLS with genome size as the independent variable and mean species protein concentration as the dependent variable. As above, statistical analyses were conducted using R v 4.6.0, and the PGLS analysis used the packages *caper* and *ape* (Orme, et al. 2011; R Core Team 2025).

## Results

### Basal cellular respiration rate and Na+/K+-ATPase pump respiratory cost are broadly similar across salamander species with different cell sizes

One-way MANOVA revealed a significant effect of species on the combined dependent variables (one-way MANOVA F(32,112) = 2.35, Pillai’s trace = 1.61, p = 0.0005). Follow-up univariate ANOVAs revealed significant differences in basal cellular respiration rate of liver tissue (F(8,35) = 3.05, p = 0.01, ω² effect size = 0.27). Mean basal cellular respiration rate of liver tissue encompassed a 1.8-fold range of values, from 1.3 *J*O_2_ (pmol O_2_/mg/s) in *P. cinereus* to 2.4 *J*O_2_ (pmol O_2_/mg/s) in *P. dunni* (Table 2). Tukey post-hoc comparisons revealed that only *P. dunni* vs. *P. cinereus* are significantly different from one another (p=0.038) (Fig. 1). The mean relative respiratory cost of the Na+/K+-ATPase in hepatic tissue showed a 1.6-fold range, from 14.9% of total respiration in *P. glutinosus* to 24% in *P. montanus*. There was no significant difference in the relative respiratory cost of the Na+/K+-ATPase among species based on univariate ANOVA (F(8,33) = 0.90, p = 0.536, ω² effect size = 0.00).

**Figure 1.**
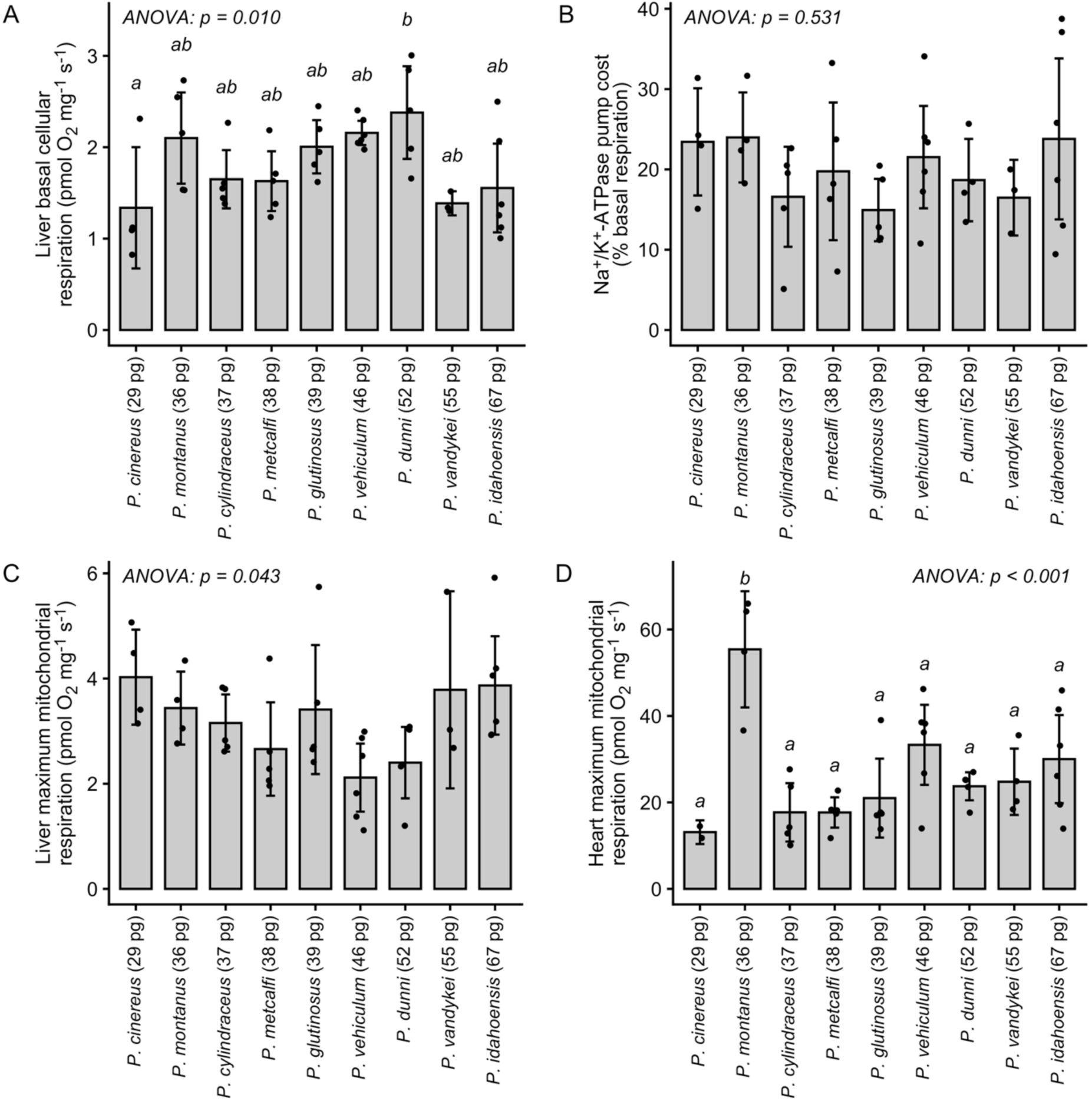
Interspecific comparisons for A) basal cellular respiration rate of liver tissue, B) relative respiratory cost of the Na+/K+-ATPase pump, C) maximal mitochondrial respiration rate of permeabilized liver tissue, and D) maximal mitochondrial respiration rate of permeabilized heart tissue. Letters denote statistically significant pairwise differences based on Tukey post-hoc tests following significant ANOVA results. Bar height shows species means, error bars represent 95% confidence intervals, and all data points are shown in black. Genome sizes are given for each species; 1 pg = 0.978 Gb.

**Table 2.** Mean protein concentrations of liver tissues.

| Species | Mean protein concentration (N) |
| --- | --- |
| *P. montanus* | 15.7% (3) |
| *P. cylindraceus* | 14.7% (3) |
| *P. metcalfi* | 12.9% (3) |
| *P. glutinosus* | 10.6% (2) |
| *P. vehiculum* | 16.9% (3) |
| *P. dunni* | 15.7% (3) |
| *P. idahoensis* | 13.1% (3) |

### Maximum mitochondrial respiratory capacity is lower and more broadly similar across species with different cell sizes in liver tissue than in heart tissue

Mean maximum OXPHOS-linked respiration rates of permeabilized liver tissue ranged from 2.1 *J*O_2_ (pmol O_2_/mg/s) in *P. vehiculum* to 4.9 *J*O_2_ (pmol O_2_/mg/s) in *P. vandykei*, with significant differences among species (ANOVA F(8,34) = 2.3, p = 0.04, ω² effect size = 0.19) (Table 2), although no individual pairwise comparisons reached significance in the Tukey post-hoc test. Mean maximum OXPHOS respiration rates of permeabilized cardiac fibers were considerably higher than those of liver, ranging from 17.7 *J*O_2_ (pmol O_2_/mg/s) in *P. cylindraceus* and *P. metcalfi* to 55.4 *J*O_2_ (pmol O_2_/mg/s) in *P. montanus* (Table 2). There were significant differences in the maximum respiration rate of permeabilized heart fibers across species (ANOVA F(8,33) = 7.15, p = 0.00002, ω² effect size = 0.54), with *P. montanus* having significantly higher cardiac mitochondrial respiration rates than all other species based on Tukey post-hoc comparisons (p < 0.02) (Fig. 1). Maximum mitochondrial respiratory capacity was not correlated between the two tissues across species.

### Genome and cell sizes are not correlated with basal cellular respiration rates, Na+/K+-ATPase pump costs, or maximum mitochondrial respiration capacity

We found no significant correlation between genome/cell size and basal cellular respiration rate (β = 0.006, p = 0.72) or the relative cost of Na+/K+-ATPase activity (β = -0.002, p = 0.99) in liver tissue (Fig. 2). We also found no significant relationships between genome/cell size and the maximum mitochondrial respiratory capacity of either liver or heart tissue across all nine species (Liver β = -0.005, p = 0.88; Heart β = 0.178, p = 0.66) (Fig. 3,4). When *P. montanus* was removed from the PGLS, we did find a significant relationship between genome/cell size and maximum mitochondrial respiratory capacity of heart tissue in the opposite direction than our prediction (p = 0.03). Pagel’s lambda showed phylogenetic signal in basal cellular respiration rate (λ = 0.7) and maximum mitochondrial respiratory capacity (λ = 0.9) in liver, but no phylogenetic signal in relative cost of Na+/K+-ATPase activity in liver (λ = 0.0) or in heart maximum mitochondrial respiratory capacity (λ = 0.0).

**Figure 2.**
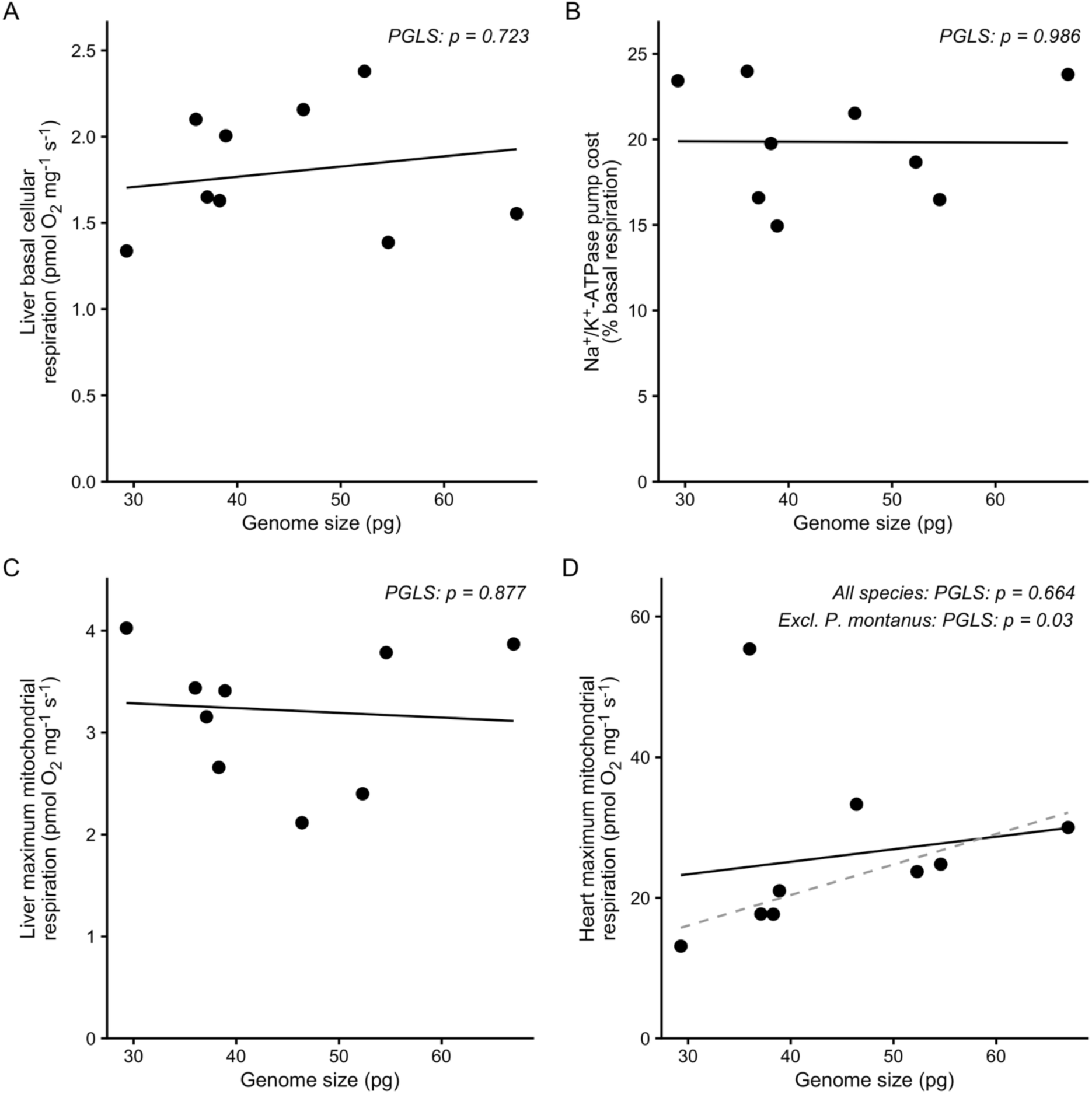
PGLS regression analyses showing no significant relationship between genome size and A) basal cellular respiration rate of liver tissue, B) relative respiratory cost of the Na+/K+-ATPase pump, C) maximal mitochondrial respiration rate of permeabilized liver tissue, and D) maximal mitochondrial respiration rate of permeabilized heart tissue. For exploratory purposes, Panel D also shows a PGLS excluding *Plethodon montanus*, which has the highest respiration rate; this exclusion results in a significant positive relationship between genome size and heart maximum mitochondrial respiration rate (dotted line).

### Protein concentration is no different across cells of different sizes

Protein concentrations of individual samples ranged from 10.2% to 18.8% (Table 2). There were no differences in protein concentration across species (ANOVA F (6,13) = 2.8, p = 0.06). There was no correlation between genome size and protein concentration based on PGLS (β = 0.0001, p = 0.89). Pagel’s lambda showed no phylogenetic signal (λ = 0.0).

## Discussion

### Larger cells do not necessarily have lower cellular respiration rates or decreased plasma membrane maintenance costs

In contrast to strong theoretical predictions, patterns present in empirical correlational studies, and direct experimental evidence in two other study stems, we find no evidence that salamander species with larger cells have lower metabolic rates at the tissue level than salamanders with smaller cells, as assessed by mass-normalized measurements of basal or maximal oxygen consumption rates *in vitro*. Contrary to our predictions, the species with the smallest cell size (*P. cinereus*) had a significantly lower basal liver cellular respiration rate than all others, and the species with a significantly higher basal cellular respiration rate (*P. dunni*) had the third largest cell size. PGLS showed no significant relationship between genome/cell size and basal respiration rate. For overall context, *Plethodon* basal liver cellular respiration values are ∼20% of the values for crocodile hepatocytes measured at 37°C (Hulbert, et al. 2002), likely reflecting differences in overall organismal metabolic rates as well as differences in assay temperature. Also in contrast to theoretical and empirical expectations, we find no evidence that salamander species with larger cells spend a smaller proportion of their total cellular respiration costs on maintenance of the membrane potential by Na+/K+-ATPase pump activity. In fact, *Plethodon* species values are not significantly different from one another, and they are also broadly consistent with measurements from other vertebrate species with more typical (i.e. much smaller) cell sizes (Milligan and McBride 1985; Hulbert and Else 2000). PGLS showed no relationship between genome/cell size and Na+/K+-ATPase cost. Finally, we find no evidence that salamander species with larger cells have lower maximum mitochondrial respiratory capacity. Liver tissue showed significant differences among species, but no pairwise comparison survived Tukey post-hoc correction, and PGLS revealed no significant relationship between genome/cell size and maximum mitochondrial respiration rate. In cardiac tissue, the species with the highest maximum mitochondrial respiratory capacity (*P. montanus*) had the second smallest cell size; this respiratory capacity was more than double that of the species with the smaller cell size (*P. cinereus*). When *P. montanus* was removed for exploratory analysis, PGLS actually revealed a positive relationship between genome/cell size and maximum mitochondrial respiration rate, in contrast to theoretical predictions. This pattern suggests that maximum mitochondrial respiratory capacity might be offsetting heart anatomical deficiencies associated with large cell size (Itgen, et al. 2022), a possibility requiring further research. Taken together, these results demonstrate that, in salamander liver and heart cells, cell size and the associated costs of maintaining ion gradients do not play a significant role in driving tissue-level metabolic costs or capacity.

What explains this departure from prediction? There are several possibilities, which are not mutually exclusive. The majority reflect a fundamental difference between comparing large and small cells from different stages of ontogeny within the same species (Jimenez, et al. 2013) or ploidy perturbation to a subset of embryos from a single clutch (Cadart, et al. 2023) (although this study also revealed the same pattern in a natural polyploid) versus comparing evolved differences in target cell size between lineages that have been diverging for tens of millions of years, as in our focal taxa.

First, contrary to geometric predictions, larger salamander liver cells may not have lower costs of membrane potential maintenance than smaller ones. This could result from one or more of the following possibilities: 1) passive diffusion rates of Na+ and K+ across the plasma membrane might not be lower in larger salamander cells than in smaller salamander cells. This condition would be met if, once cuboidal cells evolve to an extremely large target volume (as in even the smallest salamander hepatocytes), any additional increase in evolved target volume did not impact membrane potential maintenance because the intracellular distance to the plasma membrane becomes unlikely to be traversed by the ions in the center, given cytosolic viscosity. The cell would only be functionally counteracting diffusion of ions from the subset of cytosol closer to the plasma membrane to maintain ion gradients, not from the entirety of the cytosol. Analyses that image the diffusion of individual ions within the cell would allow testing of this hypothesis (Nelson, et al. 2025). 2) The enzyme efficiency of the Na+/K+-ATPase might differ across species independent of cell size, resulting in different energy costs associated with performing the work of maintaining ion gradients. In salamanders, where the metabolic rates are so low (Gatten, et al. 1992), the fitness landscape for enzyme efficiencies would be even flatter than in more metabolically expensive species, permissive to nearly neutal evolution of diverse Na+/K+-ATPase enzyme variants (Ohta 1992). This difference in enzyme efficiency could obscure any possible differences in passive ion diffusion levels across cells of different sizes. Comparing enzyme efficiencies across salamanders and other more metabolically expensive groups would allow testing of this hypothesis (Bar-Even, et al. 2011; Muir, et al. 2025).

More generally, the low metabolic rates of salamanders, and the relaxed functional constraints they impart, have left signatures at multiple levels of biological organization and may underlie the lack of correlation between cell size and respiration rate in ways beyond flattening the fitness landscape for Na+/K+-ATPase efficiency. At the DNA sequence level, salamander OXPHOS proteins show evidence of increased levels of non-synonymous substitutions, consistent with relaxed selection on OXPHOS protein function (Chong and Mueller 2013). At the biochemical level, salamanders show low OXPHOS coupling efficiency, phosphorylating relatively few ADP molecules with the proton gradient produced by the electron transport system compared to other ectothermic vertebrates; far more of the gradient is dissipated as non-specific leak, consistent with relaxed selction on ATP production (Mueller, et al. 2025). At the organ level, patterns of anatomical evolution in salamanders suggest that the structures of the heart and liver are driven by constraints imposed by large cell size on developmental systems, despite likely negative impacts on organ performance, consistent with relaxed selection on metabolic output (Itgen, et al. 2022). If OXPHOS coupling efficiency is evolving under relaxed selection in the genus *Plethodon*, then our measurements of respiration based on oxygen consumption might belie true differences in ATP use associated with differences in cell size because the same amount of oxygen consumption in different species may be associated with different amounts of ATP synthesis. Additional studies that quantify ATP use directly will address this possibility.

Linear increases in radius among smaller cell sizes will produce a more dramatic change in SA/V ratio than comparable increases in radius among larger cell sizes. Because our study system compares cell size differences among species at the largest end of the animal cell size range, the possibility exists that differences in SA/V are small enough across our focal taxa that we lacked sufficient sample sizes to detect any differences. Cadart et al. reveal high variation in oxygen consumption rate among individual tadpoles of the same ploidy level (∼1.5-fold), and used sample sizes of 11-38 tadpoles/group to detect a ∼6% decrease in mean oxygen consumption rates with increased cell size (Cadart, et al. 2023). Jimenez et al. used sample sizes of 3-10 individuals per cell size class per species to detect decreases in Na+/K+-ATPase pump costs of 4% - 85% with increased cell size, depending on species (Jimenez, et al. 2013). Although differences in cell shape of the focal cells (hepatocytes, multiciliated and goblet-like cells, and muscle cells, respectively) make exact comparisons of the magnitude of SA/V volume reductions challenging across studies, we can apply the simplifying assumption that hepatocytes and multiciliated cells are cuboidal. Estimating SA and V for our nine species of focal salamanders from the cell areas in Table 1 suggests a ∼40% reduction in SA/V ratio from our smallest- to largest-celled species. Similarly, using cell area measurements from Cadart et al. (2023), we estimate a 16% reduction in SA/V ratio between diploids and triploids. Treating muscle cells as cylindrical, we estimate that Jimenez et al included species with 17% - 79% reductions in SA/V between small and large muscle cells. Thus, our sample sizes and relative decreases in SA/V are roughly comparable to previous work. Salamanders have high levels of intraspecific variation in all respiratory variables (Figure 1), and although we consider it unlikely, we cannot formally exclude the possibility that differences exist between species, but are small enough that we did not detect them with our sample sizes.

### Tissue-level respiration rates do not correlate with whole-organism metabolic rates in Plethodon

Several studies have tested for an association between genome size and whole-organism metabolic rate among species of salamanders. Johnson et al. (2021) included five of our nine focal taxa in a larger dataset, finding no relationship between genome size and mass-specific metabolic rate measured as CO_2_ emitted at rest (Johnson, et al. 2021). There is no similarity in the rank-order between our tissue-level measurements of basal cellular respiration rates (*P. dunni* > seven *Plethodon* that are not significantly different from one another > *P. cinereus*) and Johnson et al.’s whole-organism-level measurements of metabolic rate, which were measured at 10°C and 15°C (Johnson, et al. 2021). Licht and Lowcock (1991) included two of our focal taxa, and in contrast to our tissue-level results, found that *P. cinereus* has a higher whole-organism mass-specific metabolic rate than *P. glutinosus* at 15°C and 20°C (Licht and Lowcock 1991). Discrepancies between a tissue-specific respiration rate and a whole-organism respiration rate can reflect differences in the relative size of different organs (Daan, et al. 1990; Padilla, et al. 2023); consistent with this hypothesis, among our focal species, there are 6-fold and 4-fold differences in ventricle volume and liver volume per SVL, respectively (Itgen, et al. 2022).

### Not all evolutionary changes in cell size are adaptive

Given the weight of current evidence, when does increased target cell size play a role in lowering organismal metabolic rate? The majority (13/16 species) of data comparing small and large muscle cells from single species’ ontogenies (Jimenez, et al. 2013), and two of the three developmental stages comparing diploid and triploid *Xenopus* embryos (Cadart, et al. 2023), suggest that when cell size increases and genotype is held constant (excepting gene dosage with increased ploidy), ion gradient maintenance costs decrease and metabolic rate decreases. Thus, studies inferring that decreased ion gradient costs associated with increased cell size result in lower mass-specific metabolic rates over ontogeny (geckos, ants) or in individuals inhabiting more resource-poor environnents than their conspecifics (mussels) are likely drawing the correct conclusion (Chown, et al. 2007; Starostová, et al. 2013; Dowd and Jimenez 2019). We emphasize, however, that three out of 16 species of crustaceans or fishes examined, and one out of three developmental stages in *Xenopus* examined, failed to show the predicted pattern (Jimenez, et al. 2013; Cadart, et al. 2023). Thus, decreased membrane potential cost is not an immutable structural constraint of increased cell size; some cells appear to offset geometry’s cost savings with other changes (e.g. gene expression) that remain unstudied. We caution that plasma membrane cost-savings should not be assumed, but must always be tested empirically.

What about situations in which cell size differs across distantly related species that also vary in metabolic rate? More specifically, does current evidence support the idea that large cell size played a role in the evolution of frugal life history strategies, as proposed by Szarski over 40 years ago (Szarski 1983)? We argue that the relationship between large cell size and low metabolic rate is more complex than initially envisioned, and that there are several possible scenarios that connect these two traits.

Selection for lower organism-level metabolic rate could act on numerous traits that shape overall metabolic demand, including but not limited to ion gradient maintenance costs. Other possibilities include: 1) slower protein turnover rates, which lower energy costs of translation; 2) lower proportional sizes of metabolically intensive organs; and 3) lower proportion of unsaturated fatty acids in membranes (Welle and Nair 1990; Turner, et al. 2005; White and Kearney 2013). If cell size variation is part of this initial suite of traits affected by selection for lower metabolic rate, then lower ion gradient maintenance costs would contribute to the initial stages of frugal life history evolution. However, frugal life history evolution and low metabolic rates could also initially evolve without increased cell sizes. In either case, however, the now-lower metabolic demands would relax selection against subsequent increases in cell size by decreasing diffusional limitations on oxygen and metabolites that constrain cell size. Decreases in metabolic rate driven by slower protein turnover also relax selection again subsequent increases in cell size because increased protein stability ensures that the half-life of proteins is long enough to exceed travel times across larger intracellular distances (Taylor, et al. 2024). In sum, lower metabolic rates resulting from a variety of mechanisms create a set of conditions permissive to evolutionary increases in cell size, whether or not cell size increase was part of the initial suite of traits underlying metabolic rate decrease.

Once a lineage has evolved a low metabolic rate, however, we argue that the fitness landscape for cell size becomes flattened. Any additional consequences for ion gradient maintenance costs that result from changes in cell size — if they exist at all, given other changes in enzyme efficiencies, ion diffusion dynamics, or other aspects of cell physiology — are unlikely to affect fitness in a lineage already characterized by a frugal life history strategy. Thus, cell size can evolve nearly neutrally, driven by changes in genome size that in turn reflect the accumulation of transposable element sequences. Metabolic rate itself can also evolve nearly neutrally through changes to any of its underlying drivers, leading to decoupling of cell size and metabolic rate. Lack of correlation between genome/cell size and metabolic rate across salamanders (Licht and Lowcock 1991; Gregory 2003; Johnson, et al. 2021), and high intra-specific variation in tissue-level respiration rates (Fig 1), are consistent with this scenario.

Is there anything that constrains cell size evolution or metabolic rate evolution in “frugal” lineages? The time required to move through sequences of cell differentiation and division underlying morphogenesis, during embryonic development and/or subsequent metamorphosis, imposes a constraint against larger genome and cell sizes (Gregory 2002; Liedtke, et al. 2018; Bonett, et al. 2020; Mueller, et al. 2023). Similarly, the output of developmental systems must be organ structures that meet the functional demands of the organism. Cell size increases lead to altered organ structure; lineages with low metabolic rates can tolerate more extreme changes to organ function without experiencing effects on performance or fitness, but all developmental systems presumably impose an eventual upper bound on genome/cell size (Arnold 1983; Itgen, et al. 2022).

In conclusion, we show that evolved differences in cell size can be decoupled from ion gradient maintenance costs, basal respiration rates, and maximum respiratory capacities, thus providing an explanation for the lack of correlation between genome/cell size and metabolic rate across lineages (Licht and Lowcock 1991; Gregory 2003; Uyeda, et al. 2017; Gardner, et al. 2020; Johnson, et al. 2021). Although increased cell size is expected to lower ion gradient maintenance costs when all else is held equal, other changes that accumulate in independently evolving lineages, particularly in the many traits that affect metabolic rate, can offset any impact of cell size on metabolic demand. Empirical tests that quantify Na+-/K+-ATPase activity and its contribution to cellular energy budget across diverse taxa and cell types are necessary before 1) inferring that ion gradient costs are the mechanistic connection between cell size and metabolic rate, or 2) accepting any general rules governing the evolutionary relationships between cell size, metabolic rate, and life history.

## Notes

### Competing Interest Statement

The authors have declared no competing interest.

### Funding

- U.S. National Science Foundation: 1911585
- Society for Integrative and Comparative Biology: Grants-in-aid of Research award

